# Growth temperature trigger adjustments in net photosynthesis of high-elevation plants from the Andes of Central Chile

**DOI:** 10.1101/2025.07.02.662831

**Authors:** Rodrigo Viveros, Patricia L. Sáez, Lohengrin A. Cavieres

## Abstract

Temperature significantly influences plant morphology and photosynthesis at high altitudes, shaping distinct altitudinal patterns. Lower temperatures at higher elevations typically increase leaf mass per unit area (LMA), potentially reducing mesophyll CO_2_ diffusion (gm) and chloroplast CO_2_ availability, which can limit photosynthetic rates (A_N_). However, most evidence comes from field studies where elevation-related factors, like gas pressure, water, and light, covary. Habitat type further affects plant responses, as wetland (azonal) plants may be more sensitive to thermal changes due to water’s buffering effect on temperature fluctuations. To isolate temperature effects, we conducted a controlled experiment with plants from two elevations (2600 and 3550 m a.s.l.), exposing them to two daytime temperatures (15 and 20 °C). We measured LMA, gm, and A_N_. Growth temperature significantly influenced photosynthesis and gm but not LMA. Plant origin and habitat type (zonal vs. azonal) also shaped responses. Our findings reveal that temperature impacts photosynthesis by altering mesophyll diffusion independently of LMA. Furthermore, adaptations to specific elevations and environments, such as wetlands, contribute to the variability in responses. This study highlights the nuanced role of temperature in shaping photosynthetic traits in mountain plants.

**Highlight:** Different growth temperatures affected photosynthesis independently from LMA, the response also varied from wetland species compared to non-wetland species and from plants coming from low or high elevation.

## Introduction

Decreased air temperature is one of the primary changes associated with increase in elevation (Körner, 2021). It has been suggested that this elevational decrease in temperature affects key physiological processes such as photosynthesis (Gratani et al., 2014; Kao & Chang, 2001; Kogami et al., 2001; Ma et al., 2015; Shi et al., 2015; Viveros et al., 2024). For instance, Shi et al. (2015) associated the lower leaf temperatures recorded at high elevation with lower photosynthetic rates due to low gm in evergreen trees. Similarly, reduced photosynthetic rates at high elevations were attributed to lower temperatures in *Miscanthus* species, associated with higher LMA, proposing an influence on gm (Kao and Chang 2001). Lower A_N_ in *Sesleria nitida* was also associated to lower temperatures at higher elevation in Gratani et al. (2014). Conversely, working with shrubs and grasses, Shi et al. (2015) reported higher photosynthetic rates at higher elevations, associated with higher biochemical capacity and total conductance (gs and gm). The increased A_N_ and biochemical capacity with elevation has been found in other studies as well (Fan et al., 2011; Kostopaulou and Karatassiou 2016). Nonetheless, there are other works reporting that photosynthetic rates along elevational gradients are higher at intermediate elevations due to biochemical and anatomical adaptations, but A_N_ finally decreases at the higher elevation due to low temperatures and a higher investment in survivability proteins instead of photosystems (Ma et al. 2015, Xing et al. 2024). Thus, there is no general pattern regarding changes in photosynthesis with temperature along elevations. This is likely because, with elevation, other factors such as water availability, irradiance and partial pressure of gases also change, imposing diverse effects on the photosynthetic process (Evans & Poorter, 2001; Flexas et al., 2004; Gago et al., 2020; Tosens et al., 2011). Thus, although the temperature appears to be a significant determinant of photosynthesis in high-elevation plants under field conditions, it is difficult to isolate its effects from other environmental factors that also change with elevation.

Temperature is thought to induce changes in foliar structure. For instance, Dong et al. (2020) showed increases in leaf mass area (LMA) with lower temperatures across over 700 species in Australia, similar to the results for *Quercus ilex* across 195 sites along Spain (Ogaya and Peñuelas 2007). The meta-analysis of Poorter et al. (2009) also showed that LMA tends to increase with decreasing temperatures. In the mountains of China, Zhang et al. (2020) observed increases in LMA along a gradient of around 450 m, which was attributed to decrease in temperature, given that factors such as light and relative soil moisture were similar across different elevations. Increases in LMA could affect photosynthesis in two ways: enhancing photosynthesis due to higher leaf nitrogen per area, which could positively influence key photosynthetic parameters such as maximum carboxylation rate (V_cmax_) and electron transport rate (J_max_) (Shi et al., 2015). Conversely, higher LMA could reduce photosynthesis because a higher LMA is often associated with lower g_m_ (Onoda et al., 2017), as a thicker leaf increases the diffusion distance of CO_2_ to the chloroplasts. Additionally, higher LMA could be influenced by increased in cell wall thickness, which may limit the CO_2_ diffusion in the liquid phase (Tomás et al., 2013). Recent studies have highlighted the significant role of g_m_ as a limiting factor for photosynthesis in mountain plants (e.g. Viveros et al., 2024; Xing et al., 2024). The analysis of photosynthetic limitations evaluates the balance between g_s_, g_m_ and the biochemical capacity of the leaf, determining the relative importance of each limitation, which has been observed to vary in response to different environmental conditions such as temperature, water, or nutrient availability (Niinemets et al., 2009; Gago et al., 2020). Therefore, decreases in temperature as those found along elevational gradients could reduce photosynthesis in mountain plants through increased LMA, triggering a greater mesophyll limitation.

In mountains located in mediterranean-type regions, high-elevation plants are exposed to drought, especially at the end of the growing season. However, there are azonal vegetation formations where plants do not experience drought throughout the growing season, as they growth in water saturated soils that receive water from glaciers melt runoff or groundwater outcrops. This contrasts with plants in zonal formations that grow in areas exposed to seasonal droughts. In a previous study we reported that the LMA and photosynthetic characteristics such as A_N_ and gm of zonal and azonal species responded differently to elevation (Viveros et al., 2024). Also, when comparing wetland and non-wetland plant species, Pan et al. (2020) found differences in LMA, being lower in wetland species. Further, according to the leaf economic spectrum, lower LMA could be related to vulnerability to harsh environmental conditions (Onoda et al., 2017) such as low temperatures. Additionally, azonal plants, growing in water-saturated environments, may experience greater thermal stability due to the high specific heat of water and thus could be more sensitive to changes in leaf temperature, as showed by Ruthzatz (1978), where surface soil from zonal species could reach 40°C in high Andean environments, while in wetlands the temperature never surpassed 25°C.

In the Andes of Central Chile, plants can grow up to 3600 m a.s.l. At this elevation, daytime air temperatures during growing season are around 15°C, while one kilometer downward in elevation temperatures are around 20°C (Cavieres et al., 2007). In different mountains, this altitudinal difference had been shown to be sufficient to induce changes in various foliar traits, such as LMA and photosynthetic rates (Ma et al., 2015; Shi et al., 2015; Viveros et al., 2024). However, there is no clear understanding of the factors underpinning these changes. We hypothesized that temperature drives the lower photosynthetic rates in high-elevation plants, and the mechanism underlying temperature associated variations in A_N_ involves changes in LMA and consequently in gm. Thus, when plants from high elevation are grown at higher temperatures, we hypothesize that they reduce their LMA, resulting in higher gm, lower mesophyll limitation and therefore higher A_N_. Conversely, plants from lower elevations, when grown at lower temperature, increase their LMA, leading to decreased gm, higher mesophyll limitation and hence lower A_N_. Considering the high thermal stability and lower LMA of plants growing in azonal vegetation, we also hypothesized that the effect of changes in growth temperature are bigger in azonal plants compared to zonal plants.

## Material and methods

### Experimental design

Zonal and azonal plants species growing at two elevations (2600 and 3550 m a.s.l.) in the Central Chile Andes (33.19’40’’S, 70.17’33’’ W and 33.19’34’’ S, 70.15’15’’ W) were collected and transferred to the laboratory and cultivated in a greenhouse, in pots of 500 ml, in a substrate consisting of perlite, peat, and sand in a 1:3:3 v/v. Zonal species corresponded to *Phacelia secunda* and *Bromus setifolius*, whilst azonal plants species were *Plantago barbata, Oxychloe andina*, and *Colobanthus quitensis*. Plants were watered periodically and fertilized every three weeks. After 6 weeks, zonal and azonal plants from both elevations were randomly assigned to two growth temperature regimes: 15 °C and 20 °C, with similar night-time temperature of 10 °C. These temperatures were chosen because they corresponded to the mean midday air temperatures at 10 cm aboveground during the growing season at 2600 and 3550 m a.s.l., respectively (Cavieres et al. 2007). Plants were kept at each temperature regime for at least 3 months before measurements to allow them to fully acclimate and develop new leaves in the set growth conditions.

### Chlorophyll fluorescence and gas exchange

Leaf gas exchange was measured simultaneously with chlorophyll fluorescence using a LI-6400XT gas exchange system (LI-COR Inc., Lincoln, NE, USA) with an integrated fluorescence chamber (LI-6400-40; LI-COR Inc.). Measurements were taken on a single leaf or a group of leaves, aiming to cover the entire chamber area without overlap, following the procedure described by Sáez et al. (2017). Leaf area was determined using ImageJ software (Wayne Rasband/NIH, Bethesda, MD, USA) by taking a photograph and measuring the area within the chamber. Conditions within the chamber included photosynthetically active radiation (PAR) of 1500 μmol photons m^−2^ s^−1^ for all species, except *C. quitensis*, which was set at 1000 μmol photons m^−2^ s^−1^, with 10% blue light, relative humidity between 40 and 60%, and leaf temperature fixed at 15°C and 20°C according to growth temperature. Point corrections for CO_2_ leakage were applied to all gas exchange measurements according to Flexas et al. (2007). The quantum efficiency of photosystem II (PSII) was determined according to the equation:

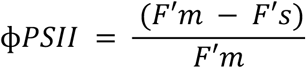

where F’s represents the steady-state fluorescence under constant light (1500 μmol m^−2^ s^−1^), and F’m represents the maximum fluorescence obtained after a saturating light pulse of 8000 μmol m^−2^ s^−1^ for 0.8 seconds. Since ϕPSII represents the number of electrons transferred per photon absorbed in PSII, the electron transport rate (ETR) can be calculated as ETR = φPSII · PPFD · αβ, where PPFD is the photosynthetic photon flux density, and αβ is a standardized parameter set at 0.45.

### Mesophyll conductance estimation

The mesophyll conductance to CO_2_ was calculated based on the combination of gas exchange and chlorophyll fluorescence, following the method described by Harley et al. (1992):

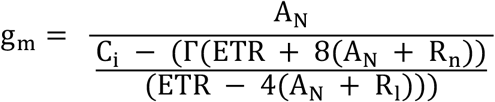

A_N_ and C_i_ were obtained from gas exchange measurements under saturating light. The rate of non-photorespiratory CO_2_ evolution under light (Rl) was assumed to be half of the dark respiration (Rn). The CO_2_ compensation point (Γ) was calculated taking into account the temperature effect over Γ according to Bernacci et al. (2002) and the altitudinal reduction of CO_2_ partial pressure (Farquhar et al., 1980). The determination of gm was used to calculate Cc.

### Leaf mass per area ratio

At each growth temperature, provenance, and species, 10 leaves were collected and photographed. Their area was measured using ImageJ software (Wayne Rasband/NIH, Bethesda, MD, USA). The leaves were then dried in an oven at 70°C for 48 hours to measure their dry mass and calculate LMA by dividing leaf mass by area.

### Quantitative analysis of photosynthetic limitations

To separate the relative control of A_N_ from the limitations imposed by stomatal conductance (gs), mesophyll conductance (gm), and biochemical capacity (lb) (gs + gm + lb = 1), the quantitative analysis of Jones et al. (1985) was used, as implemented by Grassi and Magnani (2005). The limitation of the different components was calculated as:

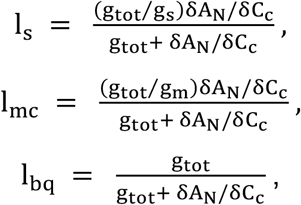

where gs is the stomatal conductance, gm is the mesophyll conductance according to Harley et al. (1992), and gtot is the total conductance for CO_2_ from the ambient air to the chloroplasts, which was calculated using the following equation:

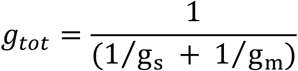

### Statistical analysis

Linear mixed-effects models were used to evaluate the effects of plant provenance (2600 and 3550 m a.s.l.) and growth temperature (15°C and 20°C) on LMA, gm, A_N_, gs, and photosynthetic limitations. In these models, species identity was included as a random factor to account for the interdependence of observations from the same species. The stomatal conductance (gs), and the stomatal and biochemical limitations (ls and lb) were log-transformed to reduce deviation from normality. Individual observations are presented untransformed to facilitate the visualization of results. Probability values were obtained from Wald’s Type II chi-square test. The normality of residuals was visually assessed using quantile-quantile plots and residual distribution plots. A Tukey post-hoc test was applied to identify specific differences between groups where there was an effect of both fixed factors and their interaction. Mixed models were conducted using the lme4 package in R statistical software (v. 4.4.1; R Core Team).

## Results

### Photosynthetic leaf gas exchange

Mean photosynthetic rates ranged from 3.3 to 7.5 μmol CO_2_ m^−2^ s^−1^ and were negatively affected by growth temperature in both zonal and azonal plants, regardless elevational provenance (Fig. 1A-B). Interestingly, gm responded differently to growth temperature in zonal and azonal plants: zonal plants did not show effects of growth temperature, whereas azonal plants exhibited lower gm values at higher growth temperatures (Fig. 1C-D). The elevational provenance effect was not significant in most variables except for gs. Mean stomatal conductance ranged from 0.08 to 0.19 mol H_2_O m^−2^ s^−1^. In azonal plants gs was only negatively affected by higher growth temperature, regardless elevation provenance, while zonal plants showed a provenance effect, with plants from higher elevation showing higher gs (Fig. 1E-F). Mean LMA ranged from 82 to 117 g m^−2^, being higher in zonal plants compared to azonal plants. However, within these groups, LMA was not significantly affected by growth temperature or plant provenance (Fig. 2A-B; Table 1).

**Table 1.**
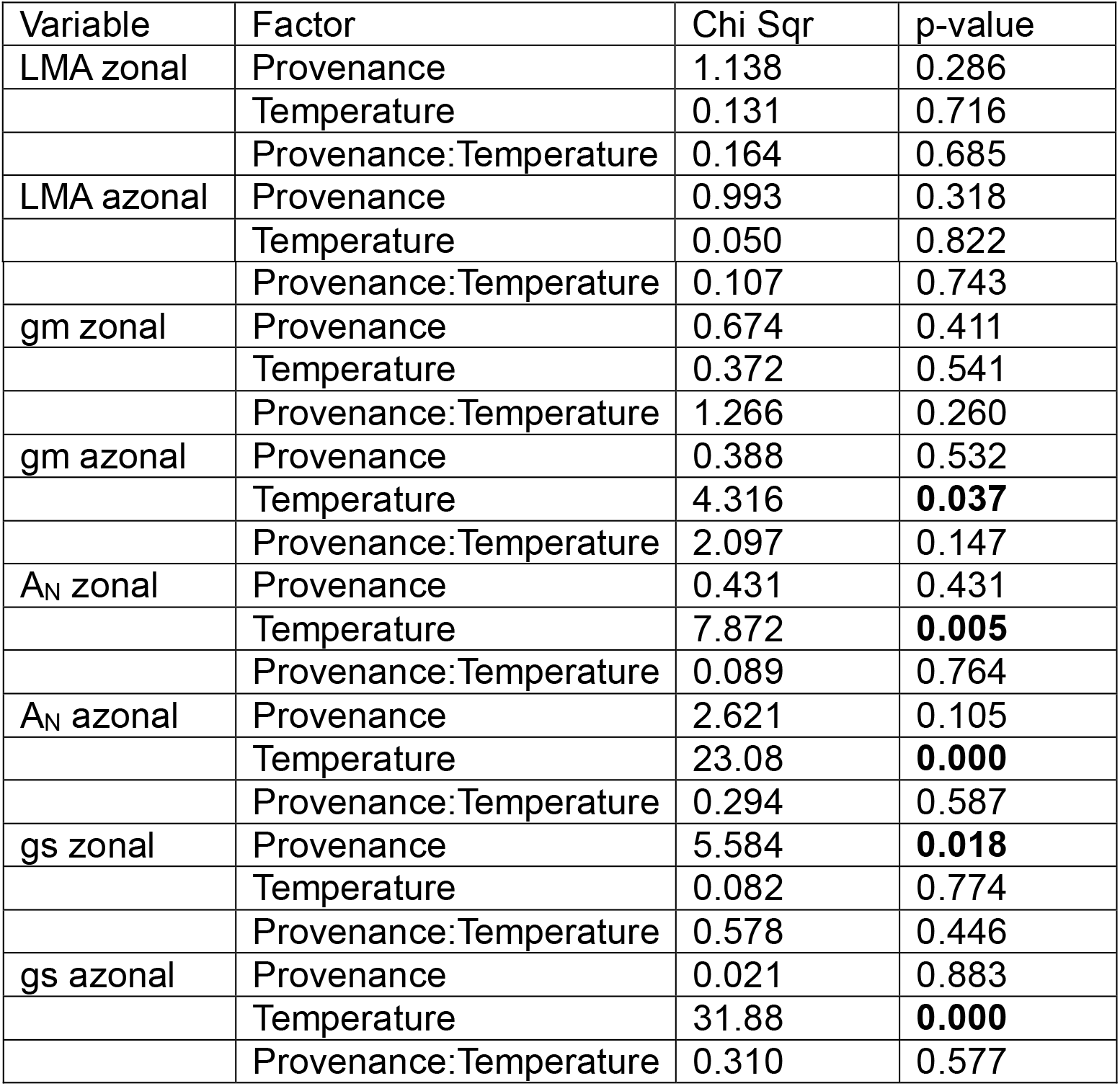
Effects of plant provenance (2600 and 3550 m a.s.l.) and growth temperature (15-20°C) on leaf structure and photosynthetic leaf gas exchange of zonal and azonal plants from the Central Andes of Chile. Significant p-values (< 0.05) from the linear mixed-effects models are presented in bold.

**Fig. 1.**
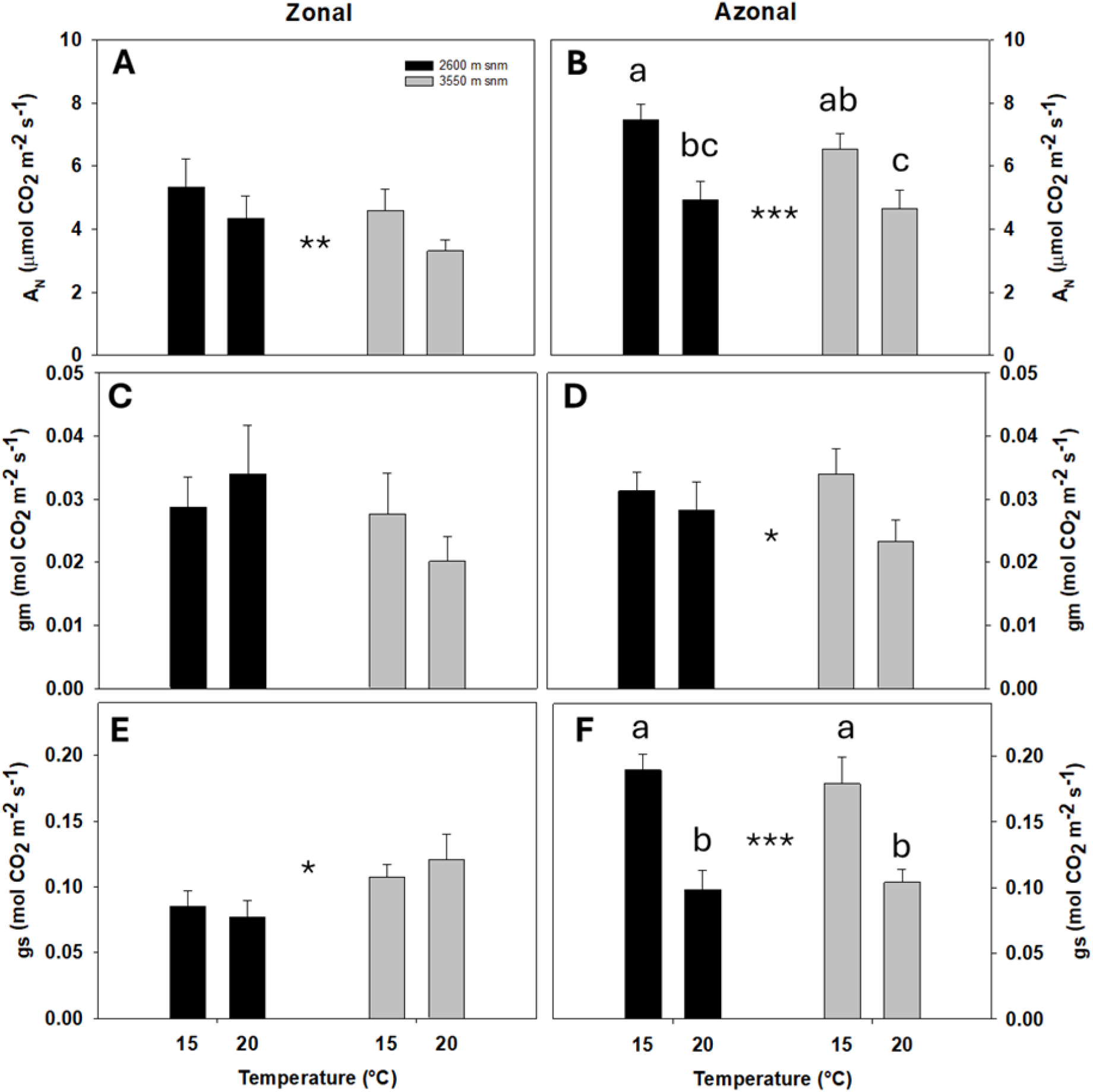
Photosynthetic response to growth temperature. Response of mean A_N_ (A-B), gm (C-D) and gs (E-F) to two growth temperatures (15 - 20°C) in zonal (left) and azonal (right) plants from two contrasting elevations, 2600 (black) and 3550 m a.s.l. (gray) n=6 for each bar, SE is presented. Asterisks indicate significant effects of temperature or provenance (p < 0.05) according to the Wald test for the linear mixed-effects models.

### Photosynthetic limitations

In zonal plants, stomatal limitation (ls) was lower in plants from higher elevations and showed a decrease when grown at 20°C (Table 2; Fig. 2A). Conversely, temperature affected the ls of azonal plants differently based on provenance, increasing with temperature in plants from 2600 m a.s.l., while no temperature effect was observed in plants from 3550 m a.s.l. (Fig. 2B). Mesophyll conductance limitation (lm) was not affected by temperature or provenance in zonal plants. In azonal plants, however, plants from different provenances responded differently to temperature; while plants from 2600 m a.s.l. showed decreased gm at higher temperature, plants from 3550 m a.s.l. did not exhibit changes in gm (Fig. 2C-D). Finally, the biochemical limitation (lb) in zonal plants increased at higher temperature. Conversely, lb in azonal plants varied with provenance, increasing in plants from 2600 m a.s.l. but remaining unchanged in plants from 3550 m a.s.l. (Fig. 2E-F).

**Fig. 2.**
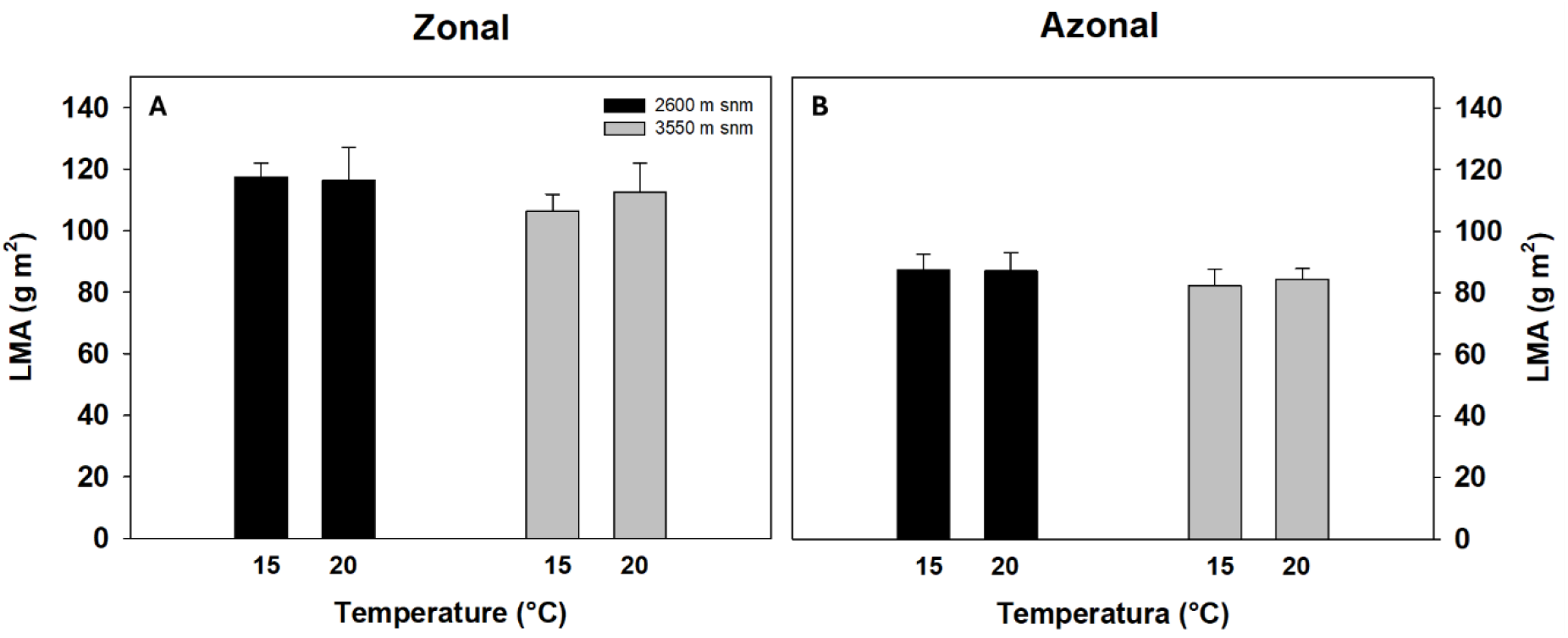
LMA response to growth temperature. Response of LMA (A-B) to two growth temperatures (15 - 20°C) in zonal (left) and azonal (right) plants from two contrasting elevations, 2600 (black) and 3550 m a.s.l. (gray). n=8 for each bar, SE is presented. Statistical differences were evaluated through Wald test for the linear mixed-effects models.

**Fig. 3.**
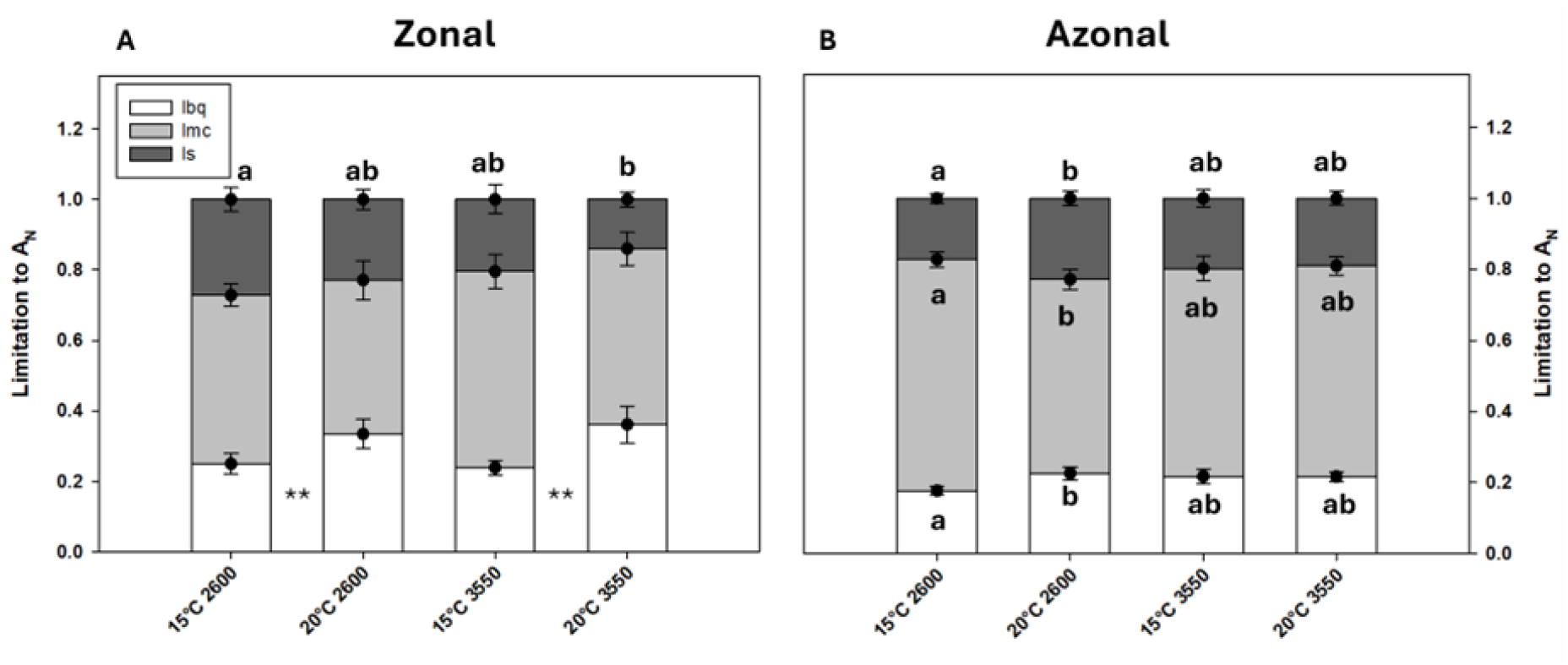
Photosynthetic limitations response to growth temperature. Mean Photosynthetic limitations: ls (A-B), lm (C-D), and lb (E-F) at two growth temperatures (15 - 20°C) in zonal (left) and azonal (right) plants from two contrasting elevations, 2600 (black) and 3550 m a.s.l. (gray). n=6 for each bar, SE is presented. Asterisks indicate significant effects of temperature or provenance (p < 0.05) according to the Wald test for the linear mixed-effects models. In cases where there is an effect of both fixed factors or their interaction, the plots show differences between means from the Tukey test (p > 0.05) in different letters when comparing the same limitation in different treatments.

**Table 2.**
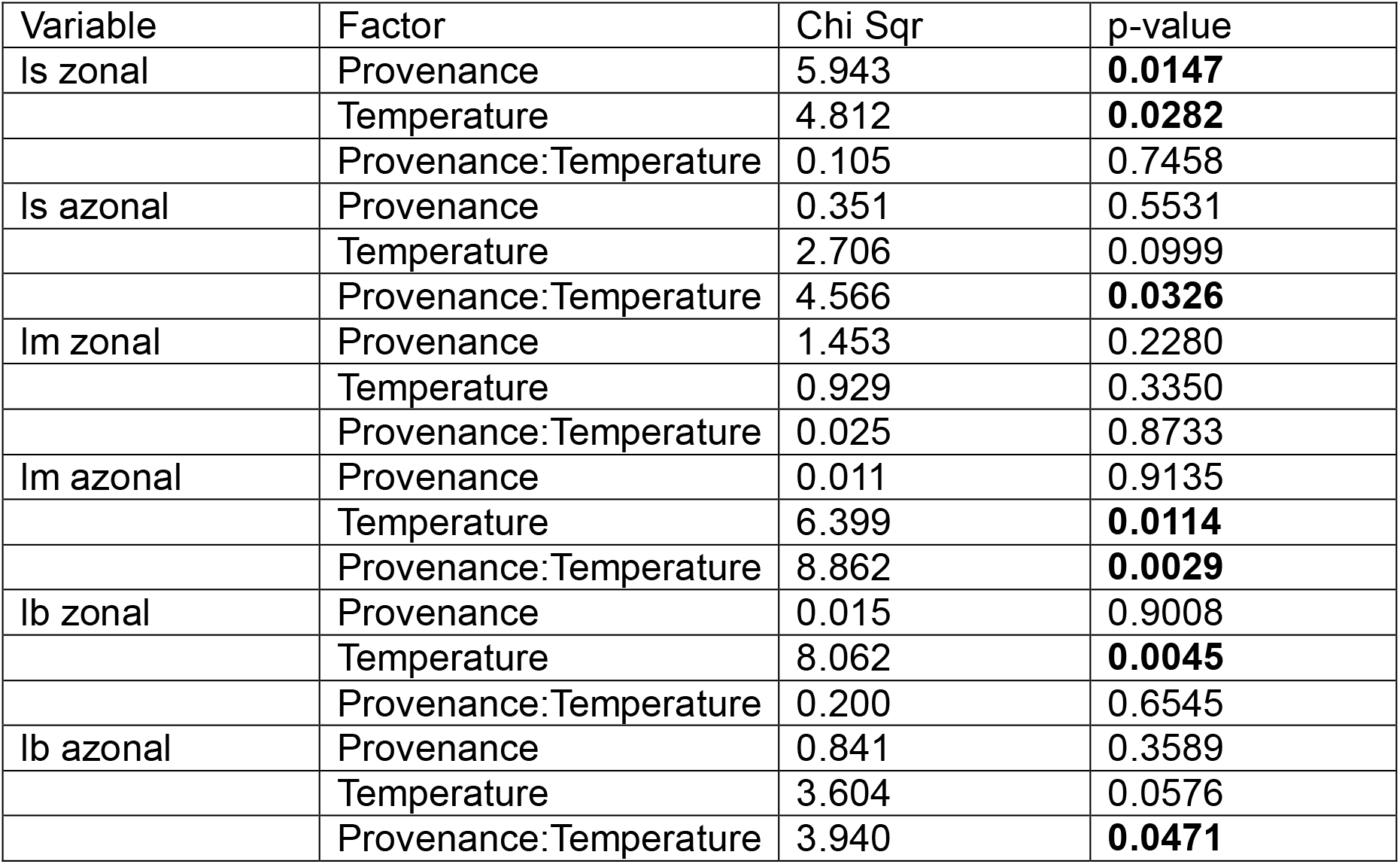
Effects of plant provenance (2600 and 3550 m a.s.l.) and growth temperature (15-20°C) on the photosynthetic limitations of zonal and azonal plants from the Central Andes of Chile. Significant p-values (< 0.05) from the linear mixed-effects models are presented in bold.

## Discussion

We hypothesized that in plants from two contrasting elevations, changes in growth temperature would lead to variation in LMA, resulting in greater limitation by mesophyll conductance (gm) and lower photosynthesis rates in plants from low elevation when grown at low temperature and the opposite for plants from higher elevation grown at high temperature. This effect was expected to be more pronounced in azonal plants due to the greater thermal stability of their environment.

According to our results, the hypothesis was rejected, as LMA did not change with different growth temperatures in both elevation provenances, contrary to observations from meta-analysis in many species (Poorter et al., 2009). In this case, the observed changes in LMA in the field could be attributed to factors such as partial pressure of gases and varying light availability (Poorter et al., 2009). However, despite similar LMA values, growth temperature exerted different effects on both gm and photosynthesis of the high-elevation species analyzed. Recent studies (Rahimi-Majd et al., 2024) support our findings by showing that, in C3 grasses, the best predictors of gm are often factors related to chloroplast arrangement and cell wall thickness, rather than LMA.

Interestingly, the effect of higher growth temperature in plant from high elevation negatively impacted the photosynthesis of the studied species. These results align with previous findings (Ma et al., 2015; Shi et al., 2015), where shrubs and grasses exhibit higher photosynthetic rates (A_N_) at higher elevations (where temperatures are lower). It is likely that at higher elevations plants may have enhanced biochemical capacities due to higher nitrogen content per area and greater Rubisco carboxylation rate (Vcmax) and electron transport rate (Jmax) (Korner & Diemer, 1987; Peng et al., 2020). This agrees with the work from Bunce et al. (2000) where cold-climate plants grown at lower temperatures deployed higher Vcmax and Jmax, and Xing et al. (2024) who found lower biochemical limitations at higher elevations. On the other hand, plants from low elevation also showed higher A_N_ at lower temperature, even when they naturally grow at higher temperatures during the growing season. This could be explained by the fact that plants present the highest photosynthetic rate at the beginning of the growing season and these values decrease as the temperature rises in the Chilean central Andes (Reyes-Bahamonde et al., 2022). Therefore, high-elevation plants may be adapted to lower growth temperatures, such as those present at the beginning of the growing season and thus decrease A_N_ when grown at higher temperatures.

As expected, zonal and azonal plants responded differently to growth temperature. While zonal plants did not show significant effects of temperature on gm or stomatal conductance (gs), azonal plants adjusted both, accompanied by decreasing A_N_. This is in line with the study of Pan et al. (2020), where plants from non-wetland areas, such as our zonal species, had higher LMA compared to wetlands plants as our azonal plants. These higher LMA were consistently associated with lower A_N_ (5.8 µmol CO_2_ m^-2^ s^-1^ in azonal plants vs. 4.3 µmol CO_2_ m^-2^ s^-1^ in zonal plants). Additionally, with higher A_N_ at low temperature, azonal plants experienced greater reductions in A_N_ when this increased (32% decrease in azonal plants vs. 23% decrease in zonal plants), which aligns with a greater sensitivity of these species to temperature changes. It is worth noting that the water in the azonal formations originates from snow melting runoff and underwater outcrops from snow and glacier melting and therefore is at low temperature. Thus, this may explain that zonal and azonal plants have different responses to temperature, with azonal plants being more sensitive to increases in growth temperature.

Another interesting finding was the effect of provenance on gs in zonal plants. Plants from 3550 m a.s.l. showed higher gs values compared to plants from 2600 m a.s.l. This effect has been observed in other studies (Gurevitch, 1992; Ma et al., 2015; Shi et al., 2015, Hernandez-Fuentes et al., 2023), being proposed that higher gs and stomatal density could act as a compensatory mechanism to cope with the lower CO_2_ partial pressure at higher elevations (Kouwenberg et al., 2007; Woodward et al., 2002). On the other hand, higher gs could naturally result from smaller cell sizes and reduced leaf expansion, as the stomata would be concentrated in smaller leaf areas, leading to higher diffusion rates per area. In any case, our results show that, regardless of temperature, zonal plants from higher elevation have higher gs values, whereas this is not the case for azonal plants, likely because these plants don’t lack water supply and hence tend to have high gs values across different elevations.

Regarding the factors limiting photosynthesis at different growth temperatures, the stomatal limitation (ls) in zonal plants was lower at higher temperature and exhibited a provenance effect, being lower in plants from higher elevation, consistent with the observed gs. In these zonal plants, biochemical limitation (lb) was not affected by provenance, but it was higher in plants grown at higher temperature. This provides further evidence of the potential adaptation of the biochemical apparatus of high-elevation plants to low temperatures, similar to the findings by Bunce et al. (2000) and proposed by Shi et al. (2015). On the other hand, the response of azonal plants was different and highly determined by plant provenance. While plants from higher elevation did not show differences in their limitations when growing at different temperatures, plants from 2600 m a.s.l. exhibited lower mesophyll limitation (lm) when grown at higher temperature, accompanied by higher lb and ls. All in all, our results revealed differential responses between zonal and azonal plants, with provenance and growth temperature affecting these responses. This highlights the importance of these factors in explaining the photosynthetic and morphological patterns observed in the field.

Overall, our results indicate that temperature may not be main responsible for the altitudinal changes in LMA but does have distinct effects on photosynthesis and its diffusive determinants, gm and gs. Additionally, the importance of the specific characteristics of plants growing at different elevations is highlighted, as these characteristics vary independently of environmental changes, such as the higher gs observed in zonal plants from higher elevation. Finally, our results underscore the differential response of zonal and azonal plants to changes in growth temperature, showing that azonal plants are more sensitive to temperature, likely due to the greater thermal stability of their water-saturated environment.

## Authors contribution

RV, LC and PS: conceptualization; RV: Formal analysis; LC and PS: Funding Acquisition; RV: Investigation; RV, LC and PS: Methodology; LC and PS: Resources; RV: writing of the original draft; RV, LC and PS: Reviewing and editing.

## References

Bernacchi, C. J., Portis, A. R., Nakano, H., Von Caemmerer, S., & Long, S. P. (2002). Temperature response of mesophyll conductance. Implications for the determination of Rubisco enzyme kinetics and for limitations to photosynthesis in vivo. Plant Physiology, 130(4), 1992–1998. 10.1104/pp.008250

Bunce, J. A. (2000). Acclimation of photosynthesis to temperature in eight cool and warm climate herbaceous C3 species: Temperature dependence of parameters of a biochemical photosynthesis model. Photosynthesis Research, 63(1), 59–67. 10.1023/A:1006325724086

Cavieres, L. A., Badano, E. I., Sierra-Almeida, A., & Molina-Montenegro, M. A. (2007). Microclimatic modifications of cushion plants and their consequences for seedling survival of native and non-native herbaceous species in the high Andes of Central Chile. Arctic, Antarctic, and Alpine Research, 39(2), 229–236. 10.1657/1523-0430(2007)39[229:MMOCPA]2.0.CO;2

Dong, N., Prentice, I. C., Wright, I. J., Evans, B. J., Togashi, H. F., Caddy-Retalic, S., McInerney, F. A., Sparrow, B., Leitch, E., & Lowe, A. J. (2020). Components of leaf-trait variation along environmental gradients. New Phytologist, 228(1), 82–94. 10.1111/nph.16558

Evans, J. R., & Poorter, H. (2001). Photosynthetic acclimation of plants to growth irradiance: The relative importance of specific leaf area and nitrogen partitioning in maximizing carbon gain. Plant, Cell and Environment, 24(8), 755–767. 10.1046/j.1365-3040.2001.00724.x

Fan, Y., Zhong, Z., & Zhang, X. (2011). Determination of photosynthetic parameters Vcmax and Jmax for a C3 plant (spring hulless barley) at two altitudes on the Tibetan Plateau. Agricultural and Forest Meteorology, 151(12), 1481–1487. 10.1016/j.agrformet.2011.06.004

Farquhar, G. D., von Caemmerer, S., & Berry, J. A. (1980). A biochemical model of photosynthetic CO2 assimilation in leaves of C3 species. Planta, 149(1), 78–90. 10.1007/BF00386231

Flexas, J., Bota, J., Loreto, F., Cornic, G., & Sharkey, T. D. (2004). Diffusive and metabolic limitations to photosynthesis under drought and salinity in C3 plants. Plant Biology, 6(3), 269–279. 10.1055/s-2004-820867

Flexas, Jaume, Diaz-Espejo, A., Galmés, J., Kaldenhoff, R., Medrano, H., & Ribas-Carbo, M. (2007). Rapid variations of mesophyll conductance in response to changes in CO 2 concentration around leaves. Plant, Cell and Environment, 30(10), 1284–1298. 10.1111/j.1365-3040.2007.01700.x

Gago, J., Daloso, D. M., Carriquí, M., Nadal, M., Morales, M., Araújo, W. L., Nunes-Nesi, A., Perera-Castro, A. V., Clemente-Moreno, M. J., & Flexas, J. (2020). The photosynthesis game is in the “inter-play”: Mechanisms underlying CO2 diffusion in leaves. Environmental and Experimental Botany, 178. 10.1016/j.envexpbot.2020.104174

Grassi, G., & Magnani, F. (2005). Stomatal, mesophyll conductance and biochemical limitations to photosynthesis as affected by drought and leaf ontogeny in ash and oak trees. Plant, Cell and Environment, 28(7), 834–849. 10.1111/j.1365-3040.2005.01333.x

Gratani, L., Crescente, M. F., D’Amato, V., Ricotta, C., Frattaroli, A. R., & Puglielli, G. (2014). Leaf traits variation in Sesleria nitida growing at different altitudes in the Central Apennines. Photosynthetica, 52(3), 386–396. 10.1007/s11099-014-0042-9

Gurevitch, J. . (1992). Differences in Photosynthetic Rate in Populations of Achillea lanulosa from Two Atitudes. British Ecological Society, 6(5), 568–574.

Harley, P. C., Loreto, F., Marco, G. Di, & Sharkey, T. D. (1992). Theoretical considerations when estimating the mesophyll conductance to CO2 flux by analysis of the response of photosynthesis to CO2. Plant Physiology, 98(4), 1429–1436. 10.1104/pp.98.4.1429

Hernández-Fuentes, C., Galmés, J., Bravo, L. A., & Cavieres, L. A. (2023). Elevation provenance affects photosynthesis and its acclimation to temperature in the high-Andes alpine herb Phacelia secunda. Plant Biology, 25(5), 793–802. 10.1111/plb.13539

Jones, H. G. (1985). Partitioning stomatal and non-stomatal limitations to photosynthesis. Plant, Cell & Environment, 8(2), 95–104. 10.1111/j.1365-3040.1985.tb01227.x

Kao, W. Y., & Chang, K. W. (2001). Altitudinal trends in photosynthetic rate and leaf characteristics of Miscanthus populations from central Taiwan. Australian Journal of Botany, 49(4), 509–514. 10.1071/BT00028

Kogami, H., Hanba, Y. T., Kibe, T., Terashima, I., & Masuzawa, T. (2001). CO2 transfer conductance, leaf structure and carbon. 529–538.

Korner, Ch., & Diemer, M. (1987). In situ Photosynthetic Responses to Light, Temperature and Carbon Dioxide in Herbaceous Plants from Low and High Altitude. Functional Ecology, 1(3), 179. 10.2307/2389420

Korner, Christian. (2021). Alpine Plant Life. Springer. 10.1007/978-3-642-98018-3 e-ISBN-13:

Kostopoulou, P., & Karatassiou, M. (2016). Photosynthetic response of bromus inermis in grasslands of different altitudes. Turkish Journal of Agriculture and Forestry, 40(4), 642–653. 10.3906/tar-1602-50

Kouwenberg, L. L. R., Kiirschner, W. M., & McElwain, J. C. (2007). Stomatal frequency change over altitudinal gradients: Prospects for paleoaltimetry. Paleoaltimetry: Geochemical and Thermodynamic Approaches, 66(October 2007), 215–241. 10.2138/rmg.2007.66.9

Ma, L., Sun, X., Kong, X., Galvan, J. V., Li, X., Yang, S., Yang, Y., Yang, Y., & Hu, X. (2015). Physiological, biochemical and proteomics analysis reveals the adaptation strategies of the alpine plant Potentilla saundersiana at altitude gradient of the Northwestern Tibetan Plateau. Journal of Proteomics, 112, 63–82. 10.1016/j.jprot.2014.08.009

Niinemets, Ü., Díaz-Espejo, A., Flexas, J., Galmés, J., & Warren, C. R. (2009). Role of mesophyll diffusion conductance in constraining potential photosynthetic productivity in the field. Journal of Experimental Botany, 60(8), 2249–2270. 10.1093/jxb/erp036

Ogaya, R., & Peñuelas, J. (2007). Leaf mass per area ratio in Quercus ilex leaves under a wide range of climatic conditions. The importance of low temperatures. Acta Oecologica, 31(2), 168–173. 10.1016/j.actao.2006.07.004

Onoda, Y., Wright, I. J., Evans, J. R., Hikosaka, K., Kitajima, K., Niinemets, Ü., Poorter, H., Tosens, T., & Westoby, M. (2017). Physiological and structural tradeoffs underlying the leaf economics spectrum. New Phytologist, 214(4), 1447–1463. 10.1111/nph.14496

Pan, Y., Cieraad, E., Armstrong, J., Armstrong, W., Clarkson, B. R., Colmer, T. D., Pedersen, O., Visser, E. J. W., Voesenek, L. A. C. J., & van Bodegom, P. M. (2020). Global patterns of the leaf economics spectrum in wetlands. Nature Communications, 11(1), 1–9. 10.1038/s41467-020-18354-3

Peng, Y., Bloomfield, K. J., & Prentice, I. C. (2020). A theory of plant function helps to explain leaf-trait and productivity responses to elevation. New Phytologist, 226(5), 1274–1284. 10.1111/nph.16447

Poorter, H., Niinemets, Ü., Poorter, L., Wright, I. J., & Villar, R. (2009). Causes and consequences of variation in leaf mass per area (LMA): A meta-analysis. New Phytologist, 182(3), 565–588. 10.1111/j.1469-8137.2009.02830.x

Rahimi-Majd, M., Leverett, A., Kromdijk, J., & Nikoloski, Z. (2024). Nonlinear models based on leaf architecture traits explain the variability of mesophyll conductance across plant species. February, 1–14. 10.1111/pce.15059

Reyes-Bahamonde, C., Piper, F. I., & Cavieres, L. A. (2022). Elevational variation of the seasonal dynamic of carbohydrate reserves in an alpine plant of Mediterranean mountains. Alpine Botany, 132(2), 315–327. 10.1007/s00035-022-00277-y

Ruthsatz, B. (1978). Las plantas en cojín de los semi-desiertos andinos del Noroeste Argentino : Su distribución local como adaptación a los factores climáticos , edáficos y antopogénicos de sus ambientes. Darwiniana, 21(2), 491–539. http://www.jstor.org/stable/23215605

Sáez, P. L., Bravo, L. A., Cavieres, L. A., Vallejos, V., Sanhueza, C., Font-Carrascosa, M., Gil-Pelegrín, E., Javier Peguero-Pina, J., & Galmés, J. (2017). Photosynthetic limitations in two Antarctic vascular plants: Importance of leaf anatomical traits and Rubisco kinetic parameters. Journal of Experimental Botany, 68(11), 2871–2883. 10.1093/jxb/erx148

Shi, Z., Haworth, M., Feng, Q., Cheng, R., & Centritto, M. (2015). Growth habit and leaf economics determine gas exchange responses to high elevation in an evergreen tree, a deciduous shrub and a herbaceous annual. AoB Plants, plv115. 10.1093/aobpla/plv115

Tomás, M., Flexas, J., Copolovici, L., Galmés, J., Hallik, L., Medrano, H., Ribas-Carbó, M., Tosens, T., Vislap, V., & Niinemets, Ü. (2013). Importance of leaf anatomy in determining mesophyll diffusion conductance to CO2 across species: Quantitative limitations and scaling up by models. Journal of Experimental Botany, 64(8), 2269–2281. 10.1093/jxb/ert086

Tosens, T., Niinemets, Ü., Vislap, V., Eichelmann, H., & Castro Díez, P. (2011). Developmental changes in mesophyll diffusion conductance and photosynthetic capacity under different light and water availabilities in Populus tremula: How structure constrains function. Plant, Cell and Environment, 35(5), 839–856. 10.1111/j.1365-3040.2011.02457.x

Viveros, R., Anic, V., Cavieres, L. A., Sáez, P. L., Ramírez, C., Labra, N., Cavieres, L. A., Fuentes, F. (2024). Mesophyll conductance limits photosynthesis and relates to anatomical traits in high-elevation plants in the Andes. Environmental and Experimental Botany, 226(May). 10.1016/j.envexpbot.2024.105916

Woodward, F. I., Lake, J. A., & Quick, W. P. (2002). Stomatal development and CO2: Ecological consequences. New Phytologist, 153(3), 477–484. 10.1046/j.0028-646X.2001.00338.x

Xing, H., Chen, J., Gong, S., Liu, S., Xu, G., Chen, M., Li, F., & Shi, Z. (2024). Variation in photosynthetic capacity of Salvia przewalskii along elevational gradients on the eastern Qinghai-Tibetan Plateau, China. Plant Physiology and Biochemistry, 212(May), 108801. 10.1016/j.plaphy.2024.108801

Zhang, L., Yang, L., & Shen, W. (2020). Dramatic altitudinal variations in leaf mass per area of two plant growth forms at extreme heights. Ecological Indicators, 110(July 2019), 105890. 10.1016/j.ecolind.2019.105890

